# Automated cell tracking using 3D nnUnet and Light Sheet Microscopy to quantify regional deformation in zebrafish

**DOI:** 10.1101/2024.11.04.621759

**Authors:** Tanveer Teranikar, Saad Saeed, The Van Le, Yoonsuk Kang, Gilberto Hernandez, Phuc Nguyen, Yichen iing, Cheng-Jen Chuong, Jin Young Lee, Hyunsuk Ko, Juhyun Lee

## Abstract

Light Sheet Microscopy (LSM) in conjunction with embryonic zebrafish, is rapidly advancing three-dimensional, *in vivo* characterization of myocardial contractility. Preclinical cardiac deformation imaging is predominantly restricted to a low-order dimensionality image space (2i) or suffers from poor reproducibility. In this regard, LSM has enabled high throughput, non-invasive 4i (3d+time) characterization of dynamic organogenesis within the transparent zebrafish model. More importantly, LSM offers cellular resolution across large imaging Field-of-Views at millisecond camera frame rates, enabling single cell localization for global cardiac deformation analysis. However, manual labeling of cells within multilayered tissue is a time-consuming task and requires substantial expertise. In this study, we applied the 3i nnU-Net with Linear Assignment Problem (LAP) framework for automated segmentation and tracking of myocardial cells. Using binarized labels from the neural network, we quantified myocardial deformation of the zebrafish ventricle across 4-6 days post fertilization (dpf). Our study offers tremendous promise for developing highly scalable and disease-specific biomechanical quantification of myocardial microstructures.

## INTRODUCTION

Investigating genetic, structural, or metabolic biophysical characteristics in response to cell dysfunction or targeted therapeutic intervention, fundamentally drives biomedical research [1–3]. A comprehensive understanding of cell-associated genomics, disease precursors or imaging biomarkers through cutting-edge biomedical technologies in the last decade, has vastly benefitted disease modeling and clinical translational of pharmaceuticals [4–6].

With respect to cardiovascular disease (CVi) in particular, correlation of cardiac cell lineage and associated phenotypes is instrumental to characterize tissue remodeling owing to hemodynamic malformations. From a biomechanics perspective, the human heart is a kinetic pump undergoing mechanical stimuli across an entire cardiac cycle [9,10]. The continuous cardiac motion of delivering oxygenated blood to organ systems is performed by cardiomyocytes (CM), experiencing anisotropic mechanical compliance based on location within the heart [11,12,13]. Consequently, identification of novel hemodynamic biomarkers is emerging a crucial research topic, to quantify local tissue deformation and cardiac output [13–15].

Understanding the complex mechanotransduction of heart tissue requires diagnostic imaging pipelines capable of multidimensional, *in vivo* cell tracking in translational animal models, thereby mimicking patient-specific heart disease. In this regard, zebrafish have emerged as robust gene-editing platforms for cardiac cell tracking or chemical screening of pharmaceuticals. This is predominantly owing to optical clarity of the specimen, low-cost maintenance, and high reproducibility of transgenesis[16–19]. Moreover, zebrafish share evolutionarily conserved gene signaling pathways with humans [7,16]. However, visualizing dynamic zebrafish heart cells is a daunting task, owing to their very small size (100-150 microns) and rapid motion (∼170 bm) [26]. Consequently, transgenic zebrafish studies have led to the development of advanced microscopy techniques that facilitated cell tracking across varying spatiotemporal scales.

In this regard, Light Sheet Microscopy (LSM) has emerged as a powerful imaging modality in zebrafish biologists, owing to cellular resolution while providing large penetration depths up to centimeter scale [7,27–29]. Moreover, LSM offers promising avenues for autonomous microscopy, in conjunction with semantic segmentation networks requiring extensive training data. This can be attributed to rapid 3i volume acquisition times, up to 30 frames per second (fps). However, current AI research is focused on segmentation of biomedical imaging data, utilizing Convolutional Neural Networks (CNNs), Support Vector Machines (SVM) or ieep Learning (iL). Hence, classification/segmentation performance is adversely affected by high data dimensionality or sparseness (less features). Moreover, current iL trends focus on manual annotation of region-of-interest (ROI) for training, in addition to non-customizable network architectures. Thereby, hindering the development of scalable segmentation models that perform generalized classification across varying imaging modalities. Other drawbacks include tradeoffs between computational resources and network depth/feature complexity.

Consequently, nnUnet has recently emerged as a highly versatile, self-adaptive framework for medical image segmentation. Unlike traditional iL methods, that require extensive manual architecture/hyperparameter tuning, nnU-Net identifies optimal network configurations based on domain-specific data [20,21]. Consequently, nnUnet has significantly reduced tradeoffs between computing power, image patch size, and GPU memory limitations, beleaguering conventional iL algorithms. In addition, instance normalization implemented in nnUnet preprocessing, enables style transfer from label images. Hence, enabling generalization of segmentation performance with respect to multimodality imaging.

In this study, we implemented a zebrafish-specific 3i nnU-Net to segment and track myocardial nuclei 4i (3i+time) trajectories acquired using LSM. While earlier studies using nnU-Net focus on 2i or 3i image segmentation, we have extended its application to higher dimensionality object tracking. In addition, we integrated Laplacian-of-Gaussian (LoG) in conjunction with morphological operators as a novel preprocessing strategy within the nnUnet framework. Consequently, we quantified significant improvements in iice coefficient, Jaccard values and time per epoch using the LoG preprocessing. Further, we quantified the area displacement of nuclei trajectories segmented using nnUnet and assessed declining area ratio trends across 4-6 days post fertilization (dpf). Thus, demonstrating the scalability and efficacy of nnUnet with respect to automated, multiscale object segmentation in higher-order image dimensionality

## MATERIALS AND METHODS

### Zebrafish lines, husbandry, and maintenance

Experiments were performed using transgenic *Tg(cmlc:GFPnuc)* zebrafish embryos obtained from spawning adult zebrafish maintained under the UT Arlington Animal Core Facility and Use Committee (IACUC) protocol (A17.014). iue to cardiac myosin light chain (cmlc) contributing majorly to the heart contractile apparatus, wild-type *Tg(cmlc:GFPnuc)* zebrafish were used to analyze differentiated cardiomyocytes. Green fluorescent protein is expressed primarily in myocardial nuclei through *cmlc* promoter, enabling wall deformation analysis by visualizing cardiomyocyte trajectories. To suppress pigmentation, 0.0025% 1-phenyl 2-thiourea (Sigma-Aldrich, St-Louis, MO) was introduced to E3 embryonic medium between 20-24 hours post fertilization [22,23]. 0.05% tricaine (MS 222, E10521, Sigma-Aldrich, St-Louis, MO) was used to sedate the embryos embedded in 0.5% low-melt agarose gel to avoid sample movement during imaging[24]. Further, agarose gel was transferred to a Fluorinated Ethylene Propylene (FEP) tube (1677L, IiEX, Chicago, IL) for sample mounting and refractive index matching between embryos and the surrounding medium (Refractive index of water = 1.33, refractive index of agarose and FEP tube = 1.34).

### Light sheet microscopy (LSM) implementation

Our home-built light sheet microscope features a single-side illumination pathway consisting of a cylindrical lens (LJ1695RM, Thorlabs) and 4x objective lens (4X Plan Apochromat Plan N, Olympus, Tokyo, Japan) to collimate cylindrical light sheet. A mechanical slit (VA100C, Thorlabs) was used to vary light sheet thickness between 1-3 um, to sample growing ventricular circumference (100-200 um) across varying developmental stages and anisotropic nuclei sizes (4-12 um). Optical sectioning of the sample was executed by a iC servo motor actuator (Z825B, Thorlabs), with z-step velocity and acceleration set at 0.005 mm/s. A single color channel detection pathway features a water dipping lens (20x/0.5 NA UMPlanFL N, Olympus, Tokyo, Japan), infinity corrected tube lens (TTL 180-A, Thorlabs), and sCMOS camera (ORCA flash 4.0, Hamamatsu, Japan). The sCMOS Camera [pixel size = 6.5 um) enabled non-gated 4i (3i + time) cardiac volume acquisition at rapid exposure times between 30–50 ms. As the zebrafish ventricle undergoes periodic deformation from peak systole to end-diastole, optical sections were captured at different time points during 4–5 cardiac cycles. In this regard, volumes were reconstructed post hoc to ensure alignment between adjacent optical sections. This process involves estimating the cardiac cycle period via least squares intensity minimization and adjusting relative period shifts to synchronize independent cardiac cycles. In a previous study, we comprehensively discussed the in silico synchronization of 4i volumes as per specific cardiac phases [25].

### 3D nnUnet implementation

The nnU-Net v2 was implemented in Python 3.10 using PyTorch v2. Batchgenerators 0.21 was used for data augmentation. Python libraries that were also used include tqdm 4.66.5, dicom2nifti 2.2.10, scikit-image 0.17.2, MedPy 0.4.0, SciPy 1.5.2, batchgenerators 0.21, NumPy 1.19.1, scikit-learn 0.23.2, SimpleITK 1.2.4 and pandas 1.1.1. nnU-Net’s source code is available on GitHub (https://github.com/MIC-iKFZ/nnUNet). Further, we implemented a novel preprocessing approach within nnUnet v2 **(Supplementary document,** https://github.com/JuhyunLeeLab/3i-Zebrafish-nnUNet.git)), focusing on the efficiency and accuracy of the inferencing obtained using 3i nnU-Net. Laplacian of Gaussian (LoG) was integrated with nnUnet v2 preprocessing framework by importing the LoG filter from the SciPy multidimensional image processing library. Edges (zero crossings) of overlapping nuclei at varying depths were localized by altering the blur operator (standard deviation) for the LoG filter. **(Supplementary Document).**

### 3D nnU-Net network configuration

4i (3i+time) LSM greyscale time series image data was acquired across 4 – 6 dpf across two cardiac cycles. Both bins for training and raw volumes were converted into 8-bit format and imported as 512 × 512 × m 8-bit arrays (raw image size). The network takes the input of size: A × A × m {A = (2^n)k | k∈N} where n is the number of levels of network, m is the image depth}. For inferencing, input patch sizes of A x A x m were generated using sliding window with half patch size overlap and output was merged using Gaussian importance weighting, Single color channel was utilized for emission, and 3×3×3 kernel sizes were utilized for convolutions during down sampling. Each computational block consisted of sequences of convolution, instance normalization, and Leaky Rectified Linear Unit (LeakyReLU) [26]. Stochastic Gradient iescent with Nesterov momentum (*μ* = 0.99) with initial learning set to 0.01 was used for network weights, in addition with combination of iice loss and Cross Entropy loss functions [26]. The default training parameter was set with 150 epochs (1 epoch = 250 iterations). To avoid overfitting, we used data augmentation methods, such as rotation, scaling, Gaussian noise, blurring, contrast and brightness adjustment, and gamma correction. Further, we obtained convolutional strides ranging from 1×2×2 to 2×2×2, based on input image size, which contributed to the network’s adaptability and performance across different image sizes. The dataset consisted of approximately 26,000 image slices. The training process consisted of 1,000 epochs (m =120). This strategy ensured efficient training for datasets across different stages of ventricular development. The output images, originally in NIfTI (.nii) format, were converted into a human-recognizable format (.tiff) for further analysis and visualization.

### Performance metrics description

We generated performance metrics, including line graphs depicting training loss, testing loss, and the iice coefficient over 1,000 epochs. iice coefficient was calculated as 2 TP / (2 TP + FP + FN), and Jaccard Index/Intersection of union was calculated as TP / (TP + FP+FN). (TP = true positive, FP = false positive, FN = false negative).

### Cell counting and trajectory tracking sz

To perform particle tracking without trajectory splitting/loss between image frames, isolating individual binarized nuclei labels from greyscale myocardial nuclei is necessary. iifference of Gaussian (ioG) bandpass filtering was used to enhance the contrast of nuclei. Briefly, n-dimensional grayscale image *I*: *T* = *T*^*n*^ is described by the function Γ_*σ*1,*σ*2_ = *I* ∗ *G*_*σ*1_ − *I* ∗ *G*_*σ*2_, produced by subtracting a Gaussian filtered image with a larger standard deviation from the same image convolved with a narrower deviation. After Otsu binarization, frame–frame linking of nuclei trajectories was performed using the linear assignment problem (LAP) framework[27]. The algorithm tracks non-branching trajectory ‘segments’ between adjacent frames through (a) Solving cost matrix for frame-to-frame linking, that takes the shape of a [n+m]x[n+m] matrix whereby n are spots in time frame t and m are spots in time frame t +1. Thereby establishing a ‘link’ or ‘no-link’ between successive trajectory segments. This step is further divided into cost function estimation across four quadrants. (b) Calculating the linking cost, where the user imposes a maximal linking distance. If the linking distance is exceeded, the cost is set to infinity. (c) Estimating cost function for non-linking, whereby the top right (n x n) and bottom left(m x m) quadrants estimated in step (a) contain costs associated with segment termination and initiation, respectively. Cell counting was performed using the 3i Object Counter plugin in ImageJ[28].

### Area ratio quantification

A triad of nuclei trajectories spaced between 5-15 um was imported into the MATLAB workspace for single area ratio quantification. In order to avoid any ambiguities for camera perspective and dynamic nuclei displacement, global coordinate vectors in x, y, and z directions are converted into local coordinates using the MATLAB function global2localcoord()[29]. Briefly, the algorithm workflow consists of calculating centroids (M x 3 vectors) of triangles (polygons) plotted between the triad of global coordinate nuclei vectors along each time step to establish the origin of local coordinate vectors at (0,0,0**) (Supplementary Figure 1).** Centroids of triangles in local coordinate space were quantified as 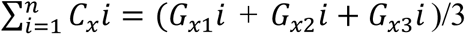, 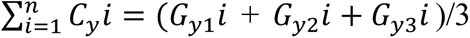, 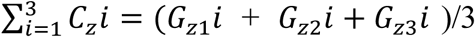, with respect to three corresponding vertices. [*G*_*x*1_*i*, *G*_*y*1_*i*, *G*_*z*1_*i*], [*G*_*X*2_*i*, *G*_*y*2_*i*, *G*_*z*2_*i*], [*G*_*z*3_*i*, *G*_*y*3_*i*, *G*_*z*3_]. For triangle ABC in local coordinate space, area was computed by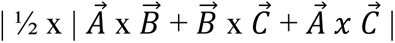[11,12].

## RESULTS

### Zebrafish CM-specific nnUnet architecture

The nnUnet is a supervised, self-configuring’ network that demonstrates robust generalization of segmentation pipeline parameters for varying datasets without any manual configuration. The nnU-Net architecture employs a U-net-based architecture, with two computational blocks per resolution stage in the encoder/decoder paths, consisting of convolution, instance normalization, and leaky RELU **(Figure 1)**. The decoder convolutional blocks mirror the encoder convolutional block, with down sampling utilizing strided (stride = 1×2×2 or 2×2×2) convolutions and up sampling implementing convolutions transposed. However, what makes nnUnet stand out is a rule-based approach that establishes interdependencies between image metadata (voxel spacing, median image size) and pipeline configuration (patch size, batch size, GPU memory constraints). This design allows the network’s fixed ‘U-net’ based architecture to iteratively optimize larger batch/patch sizes while minimizing GPU memory constraints. However, nnUnet utilizes z-score and instance normalization during preprocessing, leading to incorrect feature classification of overlapping nuclei at varying organ depth. This is due to outlier’s ‘style’ (noise) information transferred from the content image (label image) to output during normalization.

**Figure 1.**
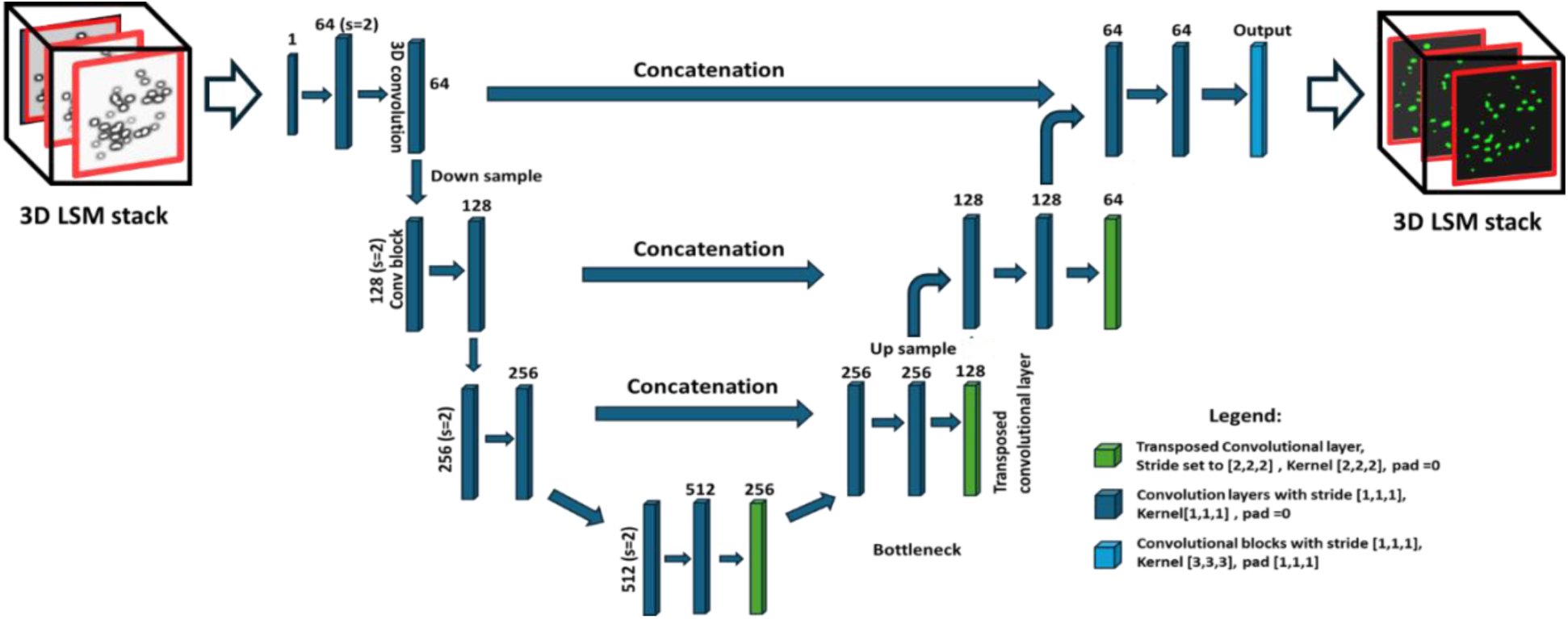
Zebrafish nnUNET architecture. Convolutional blocks within the network contain sequences of a convolution, followed by instance realization and leaky RELU activation function. Block number represents the output size, with downsampling implemented as stridden convolution and upsampling implemented as a convolution transposed.

Unlike Confocal microscopy, Light Sheet Microscopy suffers from autofluorescence along depth, saturating overlapping tissue. Hence, we integrated an additional filter-based preprocessing approach to the nnUnet framework by integrating LoG edge detection. Therefore, we were able to extract background and foreground texture information accurately across multidimensional data (gaussian blur) and isolate merged nuclei of varying scales. We validated this approach using k-fold cross-validation (k=5) by quantifying iice coefficient scores and achieved high generalization for multiscale nuclei segmentation at varying heart development stages**. (Table 1, Supplementary Video 1),**

**Table 1.** iice coefficient scores for varying zebrafish 3i+time datasets, acquired across 3 – 6 dpf (days post fertilization) with anisotropic voxel configuration, slice thickness, intensity etc.

| train<br>test | Day3 | Day4 | Day5 | Day6 |
| --- | --- | --- | --- | --- |
| Day3 | 0.8888 | 0.7381 | 0.4318 | 0.6582 |
| Day4 | 0.6621 | 0.7787 | 0.5643 | 0.6709 |
| Day5 | 0.4882 | 0.5925 | 0.8566 | 0.4874 |
| Day6 | 0.6504 | 0.6743 | 0.4547 | 0.9078 |

### Validation of zebrafish 3D nnU-Net for CM nuclei segmentation

To validate the segmentation accuracy of nnUnet for fluorescence tomographic images and its robustness (performance) with respect to localizing individual nuclei, we compared a 2i U-Net architecture[23] and 3i zebrafish specific nnUnet architecture **(Figure 2A-C).** Ground truth nuclei binarized using conventional intensity thresholding in conjunction with the watershed algorithm, were used for comparing segmentation performance with respect to 2i U-net and zebrafish specific 3i nnUnet. 2i U-Net was configured using ReLU activation and binary cross-entropy loss function with an Adam optimizer, while the 3i nnUnet relies on leaky ReLU activation and a combined iice and cross-entropy for the loss function.

**Figure 2.**
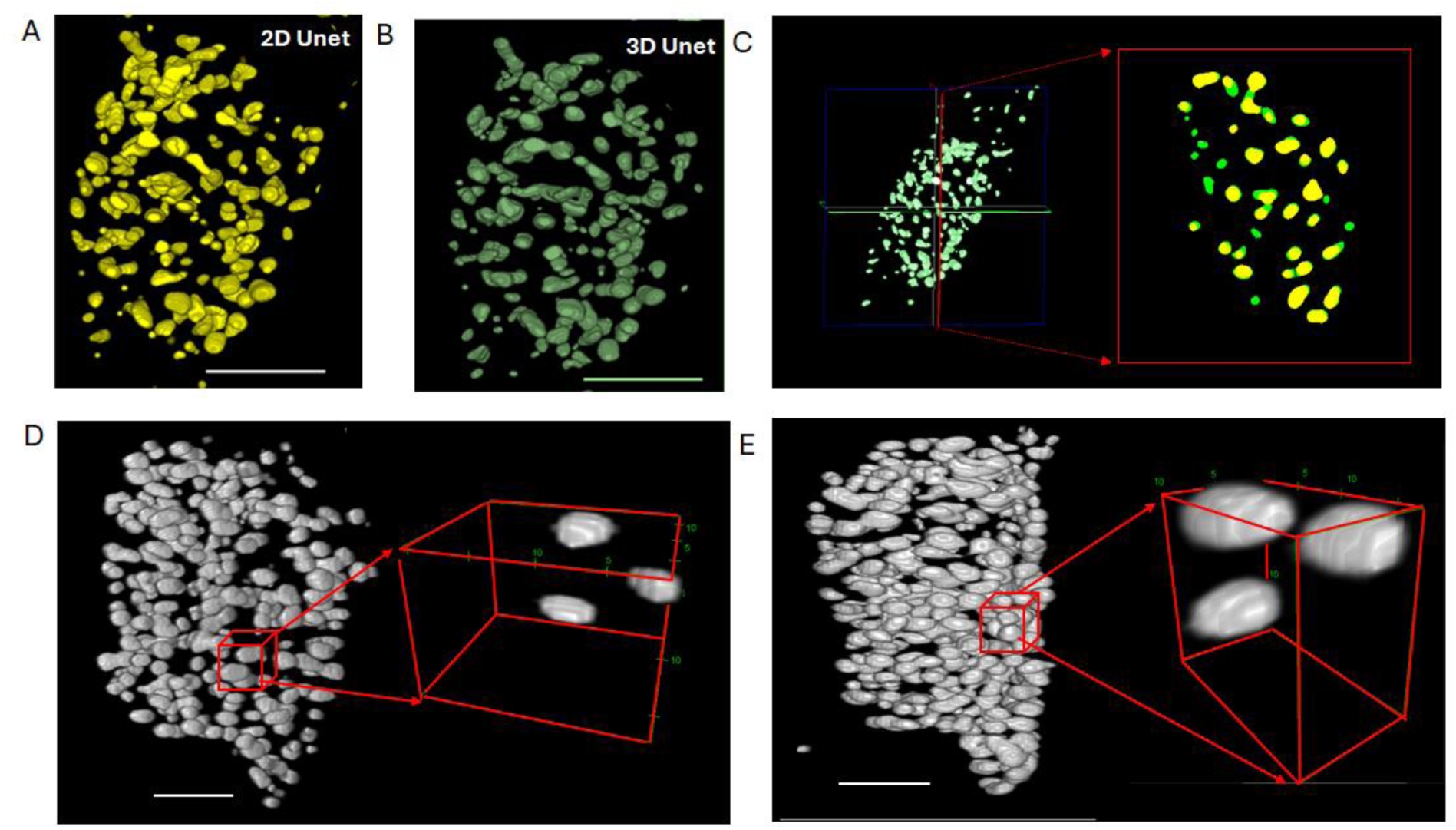
Quantifying 2D vs 3D segmentation accuracy by reconstructing 3 days post birth Tg(cmlc:GFPnuc) embryonic zebrafish ventricular myocardial nuclei. (A) Binarized nuclei volumes segmented using 2i U-Net topology with convolutional neural network (scale bar = 50 μm), (B) Binarized nuclei volumes segmented using 3i full resolution nnUnet topology (scale bar = 50 μm), (C) Cross sectional overlay of myocardial nuclei biomarkers classified using 2i Unet (yellow) and 3i nnUnet (green), indicating under segmentation by the former. (i) 5 days post fertililzation zebrafish labels produced by nnUnet preprocessed with LoG edge detector (scale bar =40 um) (E) 6 days post fertilization zebrafish labels produced by nnUnet processed with LoG edge detector scale bar = 40 um)

We quantified no significant differences in 3i nuclei volumes between ground truth nuclei datasets (164 ± 3) and 3i nnU-Net segmentation using LoG preprocessing (158 ± 4.7) **(Supplementary Figure 2A, Supplementary Video 2,3)**. On the other hand, significant under-segmentation was performed using the 2i U-Net (134 ± 5) compared to the ground truth nuclei count (**Supplementary Figure 2A).** Hence, we successfully demonstrate superior 3i network performance to automated segmentation of high dimensionality object imaging data with varying object size **(Figure 2D-2E)**.

To further validate image semantics, we performed iice coefficient and Jaccard Index comparison for the various open-source versions of nnUnet available compared to our preprocessing approach utilizing LoG edge detection. Consequently, we quantified high iice coefficients for the 3i network with 95 percent accuracy across 4 – 6 dpf nuclei segmentation, in comparison with mean 90 percent accuracy for nnUnet versions without LoG **(Supplementary Figure 3A)** for default 1000 epochs. In addition, we observed 91 percent Jaccard values for LoG based preprocessing versus 82 percent for non-LoG nnUnet versions. **(Supplementary Figure 3B)**. Hence, we successfully validated the implementation of our filter-based preprocessing approach for self-adaptive nnUnet based architecture for the segmentation of high-order dimensionality fluorescence data.

### Reconstruction of Dynamic Cell Trajectories for In Vivo Area Ratio Analysis

Using non-gated, 4i LSM, we acquired time-dependent ventricular myocardial deformation of *Tg(cmlc:GFPnuc)* zebrafish at various embryonic developmental stages **(Figure 3A).** LSM, in conjunction with 3i zebrafish nnU-Net, provides a promising end-to-end autonomous platform for imaging-based cell tracking due to its fully automated illumination and detection capability, along with self-configuring network architecture of zebrafish nnU-Net **(Figure 3B).** We integrated the multiscale imaging capability of LSM with the nnU-Net ability to handle large patch/batch sizes without affecting GPU memory constraints. Hence, we acquired and analyzed individual nuclei trajectories (15-20 microns), covering the entire ventricular circumference (50-100 microns). To reconstruct periodic deformation, we plotted nuclei labels post hoc over a complete cardiac cycle **(Figure 3C-D).** Furthermore, the nuclei pixel coordinate vectors in the global coordinate system were converted to local coordinates using a geometric center to avoid rotation and scaling artifacts affecting the scale of reconstructed objects **(Figure 3E)**.

**Figure 3.**
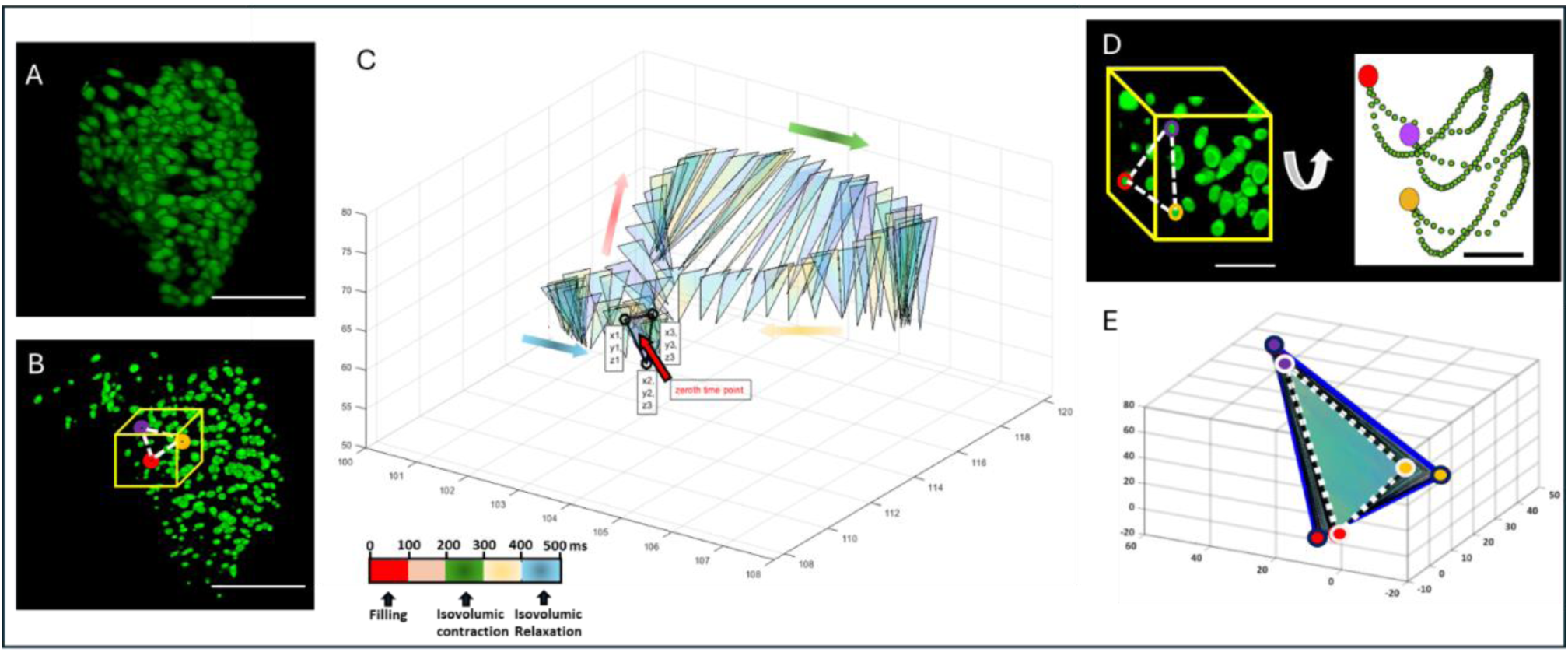
Quantifying in vivo mechanical deformation of local area ratio by tracking cardiomyocyte nuclei as markers. (A) Cardiomyocyte nuclei in a 4dpf embryonic zebrafish ventricle. (scale bar = 40 μm). (B) Binarized segmented labels produced by 3i nnuNET topology. Yellow box: Nuclei triad used for area ratio analysis, indicated by different colors (scale bar = 40 μm). (C) Plotting mechanical deformation across the cardiac cycle as a triangular vector patch between nuclei triad markers in a global coordinate system. (i) Using multi-scale imaging capability of LSM, we tracked individual nuclei trajectories across millimeter scale field-of-view. (white scale bar = 30 μm), (black scale bar = 10 μm). (E) To compare the tissue deformation, we reorient the nuclei triad from global to local coordinate transform to visualize the tissue area in the same perspective view. In area ratio computation. iotted triangle indicates default systolic cardiac phase, while solid blue boundary indicates diastolic cardiac phase in local coordinate transform.

### Zebrafish nnU-Net for Measuring Local Cardiac Tissue Deformation

Using the zebrafish 3i nnU-net, we segmented zebrafish CMs across 4-6 dpf **(Figure 4A, 5A, 6A).** We tracked the relative movements of nuclei triads in both the inner curvature of the ventricle near the inflow and the outer curvature of ventricular regions to analyze regional deformation of the ventricular myocardium across an entire cardiac cycle. **(Figures 4B-C, 5B-C, 6B-C)**. In addition, we performed 3i object counting of binarized nuclei labels produced by zebrafish specific 3i nnUnet and compared them with ground truth nuclei labels processed after LoG filtering. At 4 dpf, we quantified 257 ± 5 nuclei for the ground truth binarized labels, 264 ± 3 nuclei at 5 dpf, and 266 ± 4 nuclei at 6 dpf. For the 3i zebrafish nnUnet segmentation, the network detected 252 ± 4 nuclei at 4 dpf, 260 ± 3 nuclei at 5 dpf, and 261 ± 4 nuclei at 6 dpf. No significant differences were observed between the CM segmentation from zebrafish 3i nnUnet segmentation and ground truth at 4 dpf (<u>p = 0.15, n = 20 nuclei from 3 zebrafish hearts across two cardiac cycles</u>), 5 dpf (p = 0.43), and 6 dpf (p = 0.14) **(Supplementary Figure 2B-D)**.

**Figure 4.**
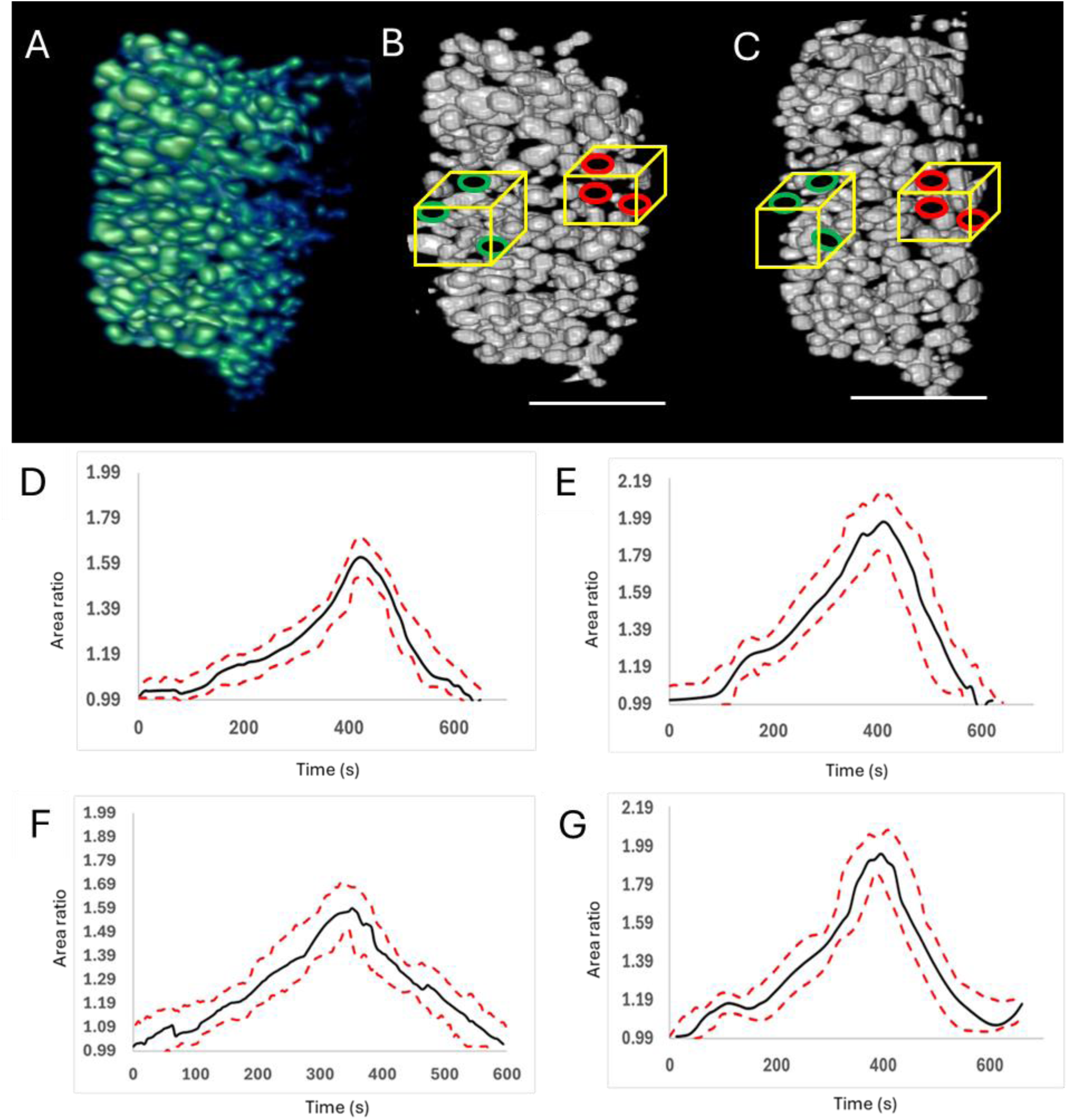
Quantifying area ratio for 4dpf zebrafish ventricular myocardium. (A) Greyscale myocardial nuclei in tg(cmlc:GFPnuc) zebrafish ventricle. (B) and (C) represent binarized nuclei labels produced by 3i zebrafish nnUNet used for area ratio quantification during cardiac diastole(The green solid boundary in the yellow box indicates nuclei in the outer curvature of the ventricle, and the red solid boundary in the yellow box indicates nuclei in the inner curvature of the ventricle), (scale bar = 50 μm). (i-E) The ground truth area ratio deformation analysis was performed in the inner and outer curvature of the ventricle, respectively, using ground truth segmentation using conventional edge detection. (F-G) The area ratio deformation analysis in the inner and outer curvature of the ventricle was performed after the segmentation from 3i nnUnet based on LoG..

**Figure 5.**
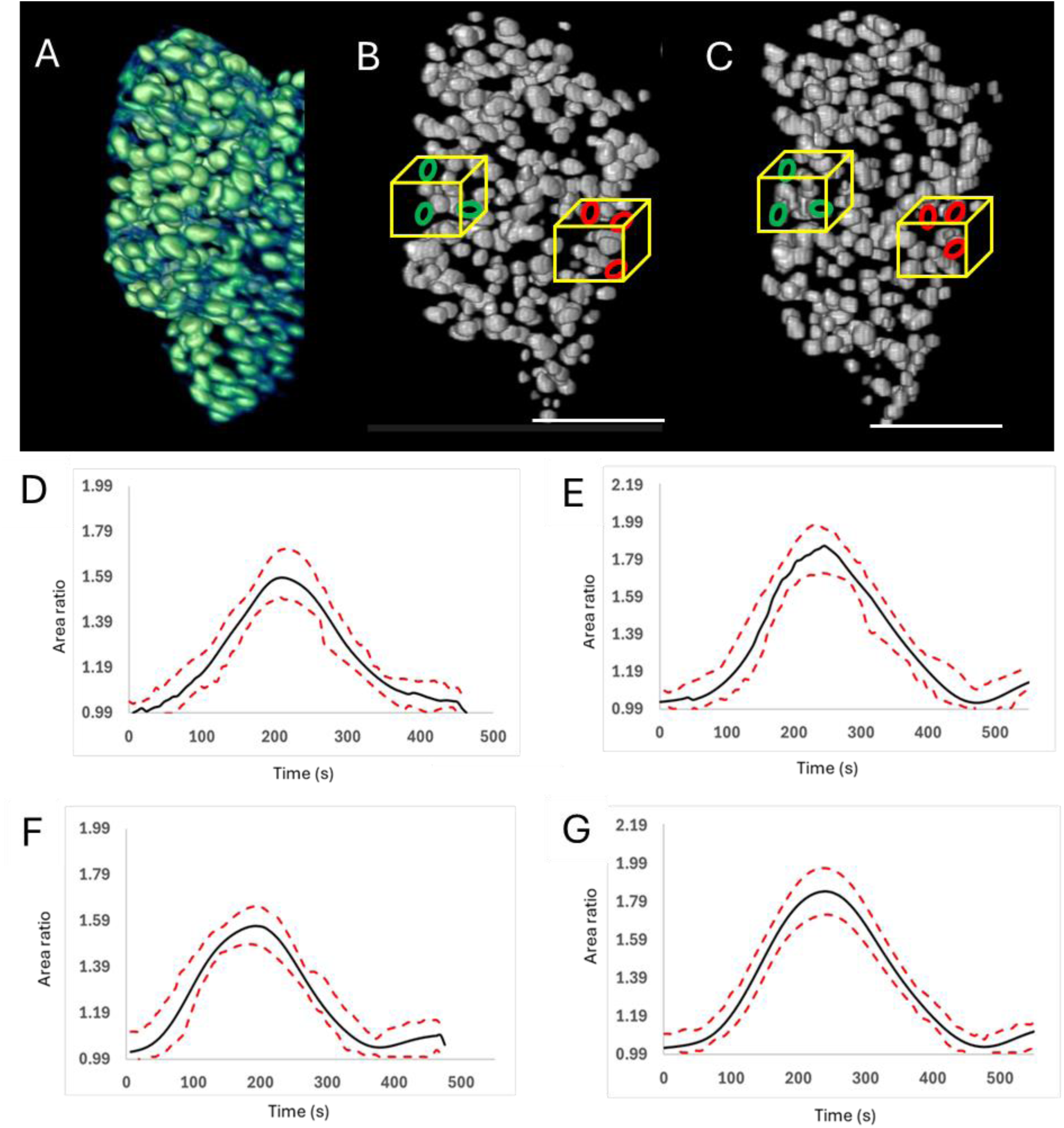
Quantifying area ratio for 5dpf zebrafish ventricular. (A) Greyscale myocardial nuclei in tg(cmlc:GFPnuc) zebrafish ventricle. (B) and (C) represent binarized nuclei labels produced by 3i zebrafish nnUNet used for area ratio quantification during cardiac diastole(The green solid boundary in the yellow box indicates nuclei in the outer curvature of the ventricle, and the red solid boundary in the yellow box indicates nuclei in the inner curvature of the ventricle), (scale bar = 50 μm). (i-E) The ground truth area ratio deformation analysis was performed in the inner and outer curvature of the ventricle, respectively, using ground truth segmentation using conventional edge detection. (F-G) The area ratio deformation analysis in the inner and outer curvature of the ventricle was performed after the segmentation from 3i nnUnet based on LoG.

**Figure 6.**
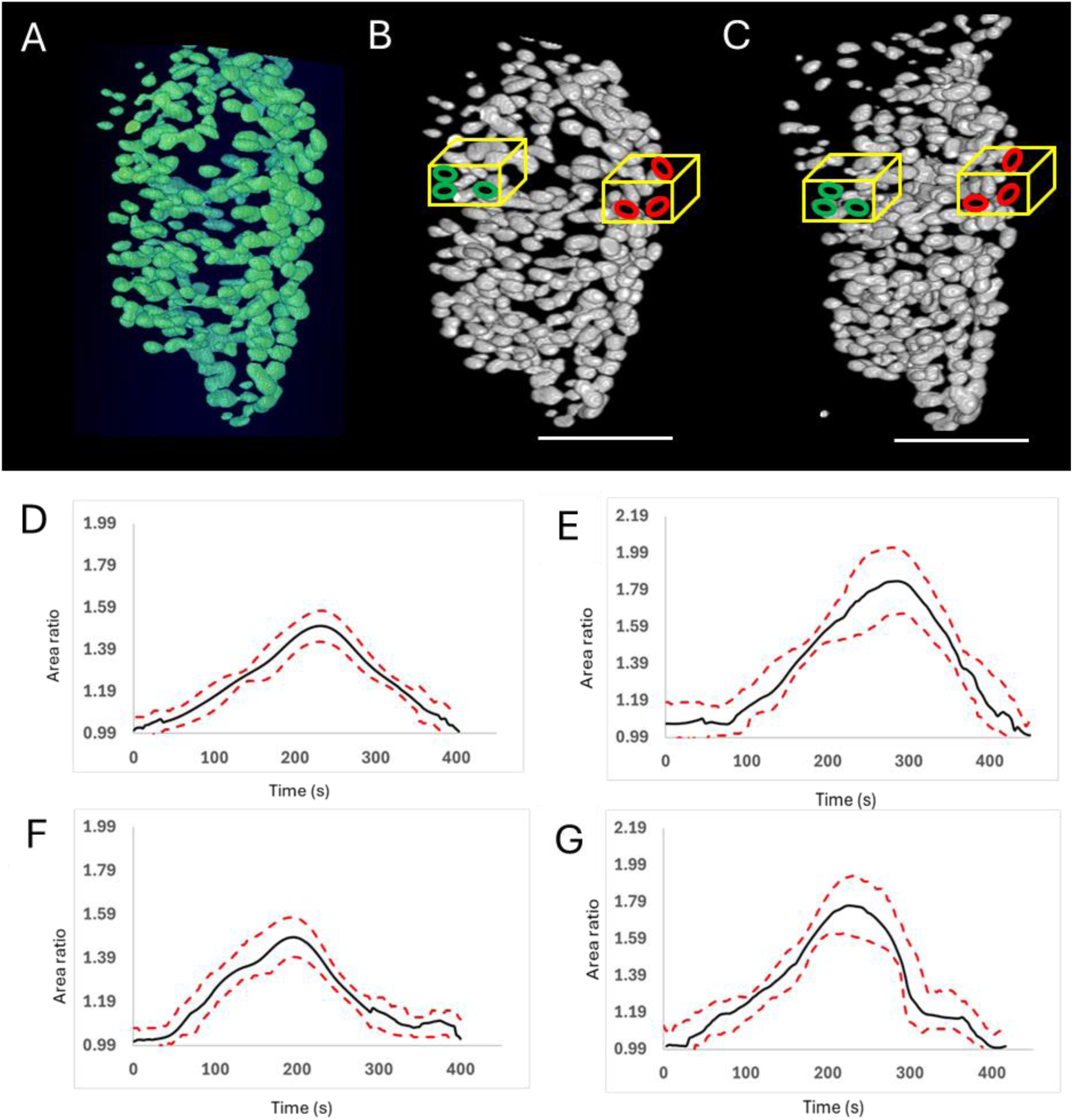
Quantifying area ratio for 6dpf zebrafish ventricular myocardium. (A) Greyscale myocardial nuclei in tg(cmlc:GFPnuc) zebrafish ventricle. (B) and (C) represent binarized nuclei labels produced by 3i zebrafish nnUNet used for area ratio quantification during cardiac diastole(The green solid boundary in the yellow box indicates nuclei in the outer curvature of the ventricle, and the red solid boundary in the yellow box indicates nuclei in the inner curvature of the ventricle), (scale bar = 50 μm). (i-E) The ground truth area ratio deformation analysis was performed in the inner and outer curvature of the ventricle, respectively, using ground truth segmentation using conventional edge detection. (F-G) The area ratio deformation analysis in the inner and outer curvature of the ventricle was performed after the segmentation from 3i nnUnet based on LoG.

We further analyzed area ratios (AR) in the inner curvature of the ventricle and found values of 1.61 ± 0.09, 1.58 ± 0.12, and 1.50 ± 0.07 at 4 dpf, 5 dpf, and 6 dpf, respectively, based on LoG ground truth segmentation **(Figures 4D,5D,6D)**. The zebrafish 3i nnU-Net showed similar ARs of 1.59 ± 0.13 (p = 0.09), 1.56 ± 0.09 (p = 0.19), and 1.48 ± 0.09 (p = 0.18) across 4-6 dpf **(Figures 4F, 5F, 6F)**. In contrast, the ground truth outer curvature of the ventricular bulge trajectories exhibited higher ARs of 1.97 ± 0.2, 1.86 ± 0.12, and 1.83 ± 0.19 at 4, 5, and 6 dpf, respectively **(Figures 4E,5E,6E).** No significant differences were observed in the 3i nnU-Net ARs for the outer curvature at 1.95 ± 0.2 (p = 0.18), 1.84 ± 0.19 (p = 0.19), and 1.81 ± 0.16 (p = 0.065) (**Figures 4G, 5G, 6G**). Thus, our segmentation pipeline successfully quantified regional myocardial wall deformation during zebrafish development, indicating that the outer curvature has higher contractility than the inner curvature.

## DISCUSSION

This study successfully validates a multiscale, cell-specific 3i+time nnUnet based architecture for quantifying local cardiac tissue deformation. By integrating a light sheet imaging, multidimensional pipeline with an automated nnUnet framework, we achieved highly reproducible insights with regards to cell trajectory tracking. In addition, we successfully demonstrated the potential of combining single particle tracking Linear Assignment Problem (LAP) algorithm[27] with self-adaptive nnU-Net[26], for generalized cell segmentation and tracking framework across varying spatiotemporal scales. Conventionally, high cell density and variable signal-to-noise (SNR) ratio have adversely affected filter-based edge detection algorithms[30]. Particularly, with respect to the classification of heterogenous tissue texture overlap in fluorescence tomography.

In this study, we integrated LoG preprocessing and watershed algorithm as part of 3i nnUnet preprocessing, to perform automated, multidimensional cell tracking for various scales of zebrafish myocardium cell maturation. Cell splitting was implemented on high-density training labels using the watershed algorithm during preprocessing, in addition to LoG edge detection for isolation of foreground and background tissue overlap. In this regard, instance normalization of nnUnet effectively transfers the ‘style’ of the training data to output. Thereby, enabling high degree of generalization with respect to cell splitting in multidimensional imaging data with variable SNR and high object density[31]. By implementing LoG edge detection and watershed algorithm in the 3i nnUnet preprocessing framework, we enhanced depth awareness of the feature extraction with respect to overlapping nuclei greyscale intensities. Consequently, we quantified no significant difference in 3i nuclei count between nnUnet preprocessing with LoG and watershed splitting, requiring several hours (68 seconds per epoch, number of epochs = 250-1000), in comparison with ground truth nuclei count produced after manual intensity thresholding over several days. This is due to instance normalization instead of batch normalization, which requires large training data.

Furthermore, we demonstrate the scalability of the LAP particle tracking algorithm in the Track Mate Fiji plugin with respect to the complexity of tracking high-order dimensionality merged objects without interpolation. The data modeling of TrackMate plugin utilizes a graph structure in contrast to using a linear data structure to store the cell trajectories[30]. Hence, cell trajectory linking is based on the strict frame–frame distance linking to better handle cell merging time-series events.

With regard to developmental biology insights, we observed anisotropic deformation in different ventricular regions, consistent with previous studies mapping regional cardiac output [11]. Significantly, no differences in Area ratios were quantified between zebrafish specific 3i nnUnet output and ground truth nuclei labels segmented using a conventional intensity thresholding approach. Hence, validating the application of nnUnet output for high-density cell tracking with variable SNR. In addition, integration of our novel preprocessing approach for 3i nnU-net, significantly improves accuracy **(Supplementary Figure 2B-D).** Previous studies suggest myocardial deformation is essential in the regulation of cardiac muscle maturation, in addition to modulation of the mechanosensitive signaling pathway such as Notch[25]. In this regard, Notch gene expression has been quantified to be significantly higher in the outer ventricular region using transgenic *tg(tp1:gfp)* zebrafish [32,33]. In this study, we have quantified significantly higher deformation in the outer ventricular region, in contrast to the atrioventricular region. Hence, future studies will focus on correlating myocardial contractility with Notch gene expression to assess the role of Notch in cardiac trabeculation.

In conclusion, our research demonstrates the efficacy of combining zebrafish specific 3i nnUnet with LSM, to achieve automated and high-precision analysis of cardiac tissue deformation in a zebrafish model. The integration of our novel preprocessing approach for 3i nnUnet, offers a powerful platform for high-throughput trajectory analysis of cell merging/division events, thereby facilitating fluorescence tomographic studies probing cellular dynamics.

## CONCLUSION

Integrating nnU-Net with LSM, enabled automated multidimensional classification and tracking of dynamic cells acquired *in vivo.* Furthermore, the integration of Laplacian of Gaussian in addition with watershed algorithm for 3i nnU-Net training, helped us achieve high-throughput trajectory localization for cell merging events, without interpolation. Consequently, our imaging workflow enabled 3d+time nuclei tracking for quantification of zebrafish myocardial strain across entire cardiac morphogenesis

## ACKNOWLEDGEMENTS

Authors would like to express gratitude to ir Yichen iing for providing embryonic zebrafish datasets for area ratio quantification. Furthermore, we are sincerely grateful to the National Institute of Health (NIH R35GM150947) and the National Research Foundation of Korea (NRF RS-2024-00419286) for supporting our work.

