## Supplementary Document for "Automated cell tracking using 3D nnUnet and Light Sheet Microscopy to quantify regional deformation in zebrafish"

### SUMMARY

The supplementary document describes a comprehensive segmentation workflow utilizing nnUnet, based on unique features of a dataset (image size, voxel data etc). The segmentation pipeline includes data preparation, model setup, training, result processing, in addition to combining multiple models (2D, 3D modes). After executing nnU-Net, the resulting model(s) can be utilized for inference on test data.

#### Setting up the environment

1. Create new environment.
    a. Enter command: conda create -n NAME python==3.10
        i. NAME is desired environment name. Ex: nn_UNet
        ii. conda create -n nn_UNet python==3.10

2. Activate environment:
    a. conda activate NAME
        i. Ex: conda activate nn_UNet
            1. Check version. Enter command: python –version

3. Cd to the directory where all the zebrafish repositories will be stored.

4. Enter command **git clone https://github.com/nasyxx/zebrafish_seg.git**
    a. If git is not installed then – **sudo apt install git – all**

b. Ex: **conda install pytorch torchvision torchaudio pytorch-cuda=11.8 -c pytorch -c nvidia**

5. Transfer the tiff_converter.py file into this directory.

6. cd into *data* directory

c. Enter command: **mkdir raw ori segmented results preprocessed cropped inference**

7. Save your environment variables.

a. Enter command: **vim ~/.bashrc**

i. If not installed enter following command: **sudo apt install vim**

1. Retry command after installation is finished

ii. If blank, double check spelling and retry.

iii. Press Insert key to begin editing.

b. Move indicator to very bottom of page and add:

**export nnUNet_raw_data_base=data/raw/**

**export nnUNet_raw=data/raw/nnUNet_raw_data**

**export nnUNet_results=data/results/**

**export nnUNet_preprocessed=data/preprocessed/**

**export**

**RESULTS_FOLDER=/home/USER/nnUNetFrame/zebrafish_seg/data/results/**

c. Press Escape key to return to command mode.

1. Enter **:x** to save and exit (Alternatively enter **:wq**)
2. Enter **:q!** to exit WITHOUT saving

8. Source your environmental variables:

d. Enter command: **source ~/.bashrc**

e. Reactivate your environment: **conda activate NAME**

i. Ex: **conda activate nn_UNet**

9. Install nnUNet repository

d. **git clone https://github.com/MIC-DKFZ/nnUNet.git**

e. **cd nnUNet**

f. **pip install -e .**

g. **pip install --upgrade git+https://github.com/FabianIsensee/hiddenlayer.git**

10. Install pytorch **https://pytorch.org/get-started/locally/**

h. Follow directions to install according to computer specifications

11. Install pdm and start project

i. **python -m pip install pdm**

j. **pdm init**

Follow the prompts. You will see:

*"Please enter the Python interpreter to use*

*0. /home/leelab/anaconda3/envs/zebrafish_seg/bin/python (3.10)*

*1. /home/leelab/anaconda3/envs/zebrafish_seg/bin/python3.10 (3.10)*

*2. /usr/bin/python3.10 (3.10)*

*Please select (0): "*

Enter whichever is your preference. Ex: *Please select (0):* 2

*"Is the project a library that is installable?*

*If yes, we will need to ask a few more questions to include the project name and build backend*

*[y/n] (n):"*

Enter: **y**

Project name: **Zebrafish Segmentation**

Project version (0.1.0): **0.1.0**

Project description (): **Auto-segmentation tool**

*"Which build backend to use?*

*0. pdm-backend*

*1. Setuptools*

*2. flit-core*

*3. Hatchling "*

Enter: **0**

"*License(SPDX name) (MIT):* **AFL-1.1**

*Author name ():* **JL**

*Python requires('*' to allow any) (>=3.10):* **>=3.10**"

12. Install remaining dependencies:

    k. **pdm run pip install tqdm**

    l. **pdm run pip install pathlib**

    m. **pdm run pip install rich**

    n. **pdm run pip install smile_config**

    o. **pdm run pip install tifffile**

    p. **pdm run pip install nnunet**

    q. **pdm run pip install nnunetv2**

        i. Note, some may not install. Keep an eye on errors for future commands to see which libraries are missing.

#### Dataset Conversion

13. Cd to *src* folder, then open dc.py

    f. Either with command: **vim dc.py**

        i. Press Insert key to begin editing

        ii. If not installed enter following command: **sudo apt install vim**

            1. Try again when installed

    g. Or may be opened and edited using Notepad

Edit <u>line 75</u>:

h. **Space = tuple(map(float,conf.space.split(“,”)))**

14. Comment out <u>lines 88, 100-119</u> using #

    i. If on vim, press Escape key to return to command mode

        i. Use command **:x** or **:wq** to save and exit, note to enter commands the command must begin with colon key “ **:** ”

        ii. Use command **:q!** to exit WITHOUT saving

    j. If on Notepad, simply save the file before exiting.

        i. Either CTRL + S

        ii. On menu bar File>Save

15. Return to directory containing zebrafish_seg repositories from github

    k. Enter in terminal while in *src* directory: **cd ..**

16. Enter command: **pdm run python -m src.dc --tod --ori path/to/raw --seg path/to/seg --database path/to/database_directory --name NAME –t XXX**

    l. Note:

        i. If environment variables are set up already, you do not need to specify --ori, --seg, --database.

        ii. path/to/raw is the pathway to the ori directory. Default: data/ori

        iii. path/to/seg is the pathway to the segmented directory. Default: data/segmented

iv. path/to/database_directory is pathway to zebrafish database. Default: data/

v. NAME is to be replaced with desired name. Ex: zebrafish_segv2

vi. XXX is to be replaced with three digit task identifier number. Ex: 777

17. Convert Task (nnUNetv1 format) to Dataset (nnUNetv2 format):

    m. **pdm run nnUNetv2_convert_old_nnUNet_dataset Path/To/Task/Folder DatasetXXX_NAME**

        i. Path/To/Task/Folder is the pathway to where the task is located. Ex: data/raw/nnUNet_raw_data/Task777_zebrafish_segv2

        ii. Replace XXX with three digit dataset identification number. Ex: 777

        iii. Replace NAME with desired name for the dataset. Ex: zebrafish_segv2

            1. Final product example: Dataset777_zebrafish_segv2

#### Plan and Preprocess

1. Edit default preprocessor to include Laplacian of Gaussian.

    *a.* Download default_preprocessor.py file containing the pre-added code. Move to directory containing the nnUNet repositories. Default preprocessor can be found at: *nnUNet > nnUNetv2 > preprocessing > preprocessors > default_preprocessor.py*

i. Delete default_preprocessor.py and replace with the updated default_preprocessor.py file. The new version will include the Lapacian of Gaussian code using scipy library.

2. Begin plan and preprocessing.
   a. Enter code:

      **pdm run nnUNetv2_plan_and_preprocess -d XXX -c 3d_fullres --verify_dataset_integrity**

   b. Replace XXX with Dataset three digit ID. Ex: pdm run nnUNetv2_plan_and_preprocess -d 777 -c 3d_fullres --verify_dataset_integrity

3. (Optional) To check to see if the new preprocessor is working as intended, utilize the tiff_converter.py file.
   a. Make new directory in data that will contain the preprocessed .tif files.
      i. (Optional) Create subdirectory for the converted .npz to .tif files for organizational purposes.
   b. Enter command: **vim tiff_converter.py**
   c. Replace input_folder = ‘path/to/preprocessed/images’
      i. Ex. input_folder = '/home/senior/nnUNetFrame/zebrafish_seg/data/preprocessed/Dataset444_zebrafish/nnUNetPlans_3d_fullres/'
   d. Replace output_folder= ’path/to/tiff/output’

i. Ex: output_folder = '/home/senior/nnUNetFrame/zebrafish_seg/data/tiff_files/Dataset444_zebrafish_seg'

e. Enter command: **pdm run python tiff_converter.py**

4. Once completed, begin training.
   a. Enter code**: nnUNetv2_train DATASET_NAME_OR_ID 3d_fullres 5**
      i. Replace DATASET_NAME_OR_ID with dataset id used for plan and preprocessing. Ex: pdm run nnUNetv2_train 777 3d_fullres 5
      ii. Results will be found in data/results
