## Supplementary Figures for "Automated cell tracking using 3D nnUnet and Light Sheet Microscopy to quantify regional deformation in zebrafish"

**Supplementary information list**

**Supplementary Figure 1**: Quantifying area ratio using cardiomyocyte trajectories acquired in vivo

**Supplementary Figure 2:** Validation for nnUnet segmentation

**Supplementary Figure 3:** Validation for 3D object counting between nnUnet and conventional intensity thresholding

**Supplementary Video 1:** Reconstructing myocardial cardiomyocyte trajectories using linear assignment problem framework

**Supplementary Video 2:** 3 days post birth zebrafish myocardial nuclei binarized using Laplacian of gaussian edge detection filter and morphological operations

**Supplementary Video 3:** 3 days post birth zebrafish myocardial nuclei segmented using 3d nnUnet and visualized across entire cardiac cycle

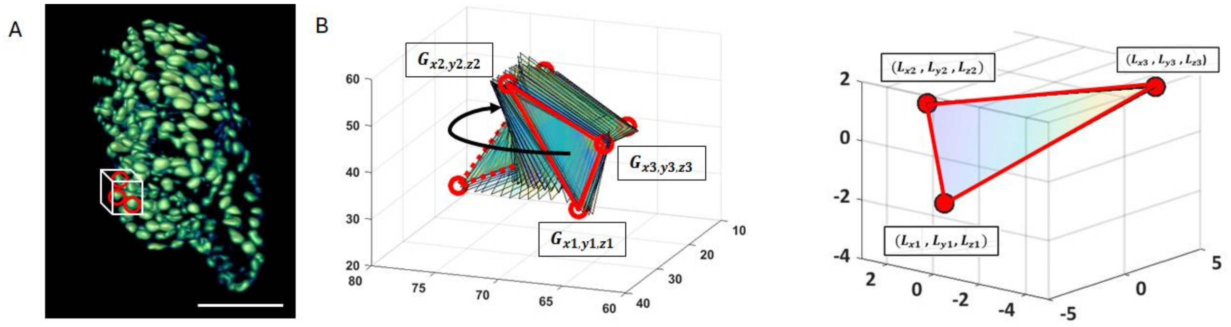


**Supplementary Figure 1. Quantifying area ratio using cardiomyocyte trajectories acquired in vivo**, (A) 6dpf myocardial nuclei in zebrafish ventricle. A nuclei triad is required to compute area of triangle for mechanical deformation (scale bar = 50 μm). (B) Triangle (polygon) plotting was performed in a global coordinate system across the entire cardiac cycle to quantify centroids required for origin. Solid red boundary indicates systole, dotted red boundary indicates diastole. (C) Nuclei position vectors reconstructed in local coordinate space for a single image frame, centered at (0,0,0) to avoid inaccuracies in area quantification due to camera perspective and nuclei displacement.

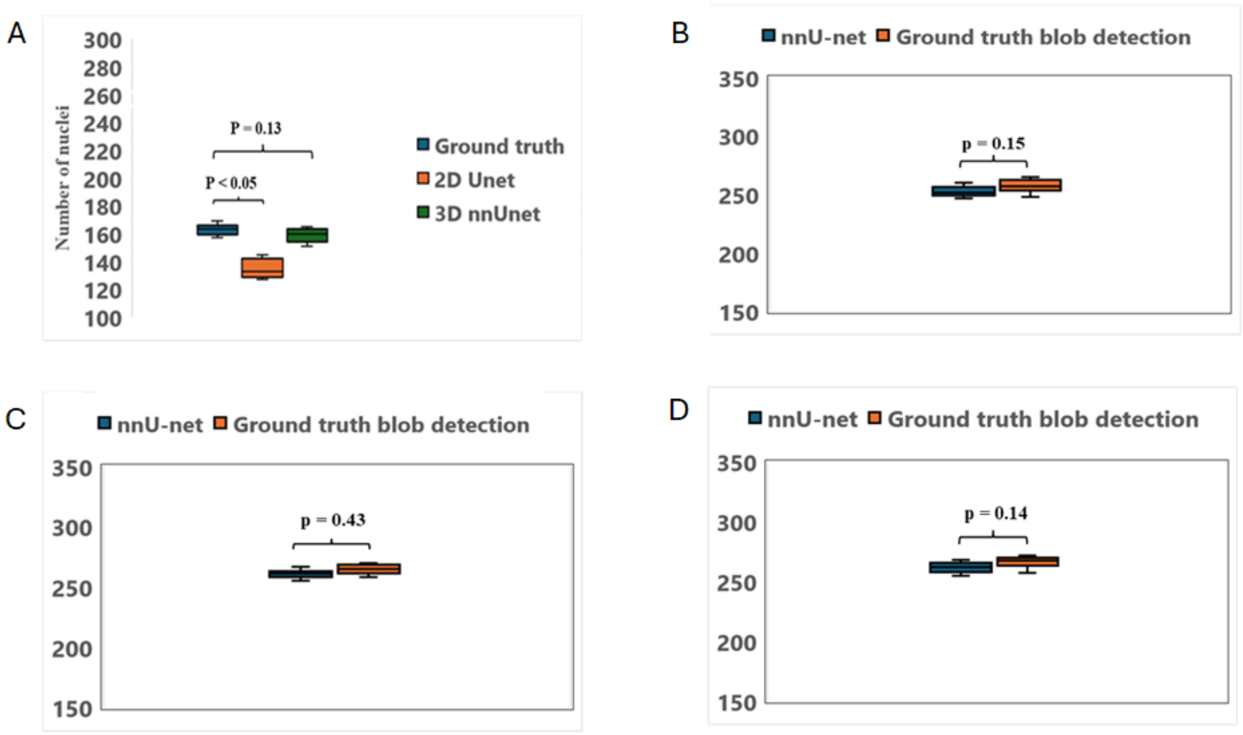


**Supplementary Figure 2. Validation for 3D object counting between nnUnet and conventional intensity thresholding segmentation** (A) T test comparing total number of nuclei segmented by 2D Unet architecture versus 3D nnUnet self-configuring network (n=20 nuclei sampled across 3 zebrafish, p value =< 0.05) (B) Nuclei count for ground truth nuclei segmentation using filter-based edge detection (Laplacian of Gaussian), in comparison with 3D nnUnet nuclei segmentation for 4 dpf dataset. (C) Nuclei count for ground truth nuclei segmentation using filter based edge detection (Laplacian of Gaussian), in comparison with 3D nnUnet nuclei segmentation for 5 dpf dataset (D) Nuclei count for ground truth nuclei segmentation using filter based edge detection (Laplacian of Gaussian), in comparison with 3D nnUnet nuclei segmentation for 6 dpf dataset

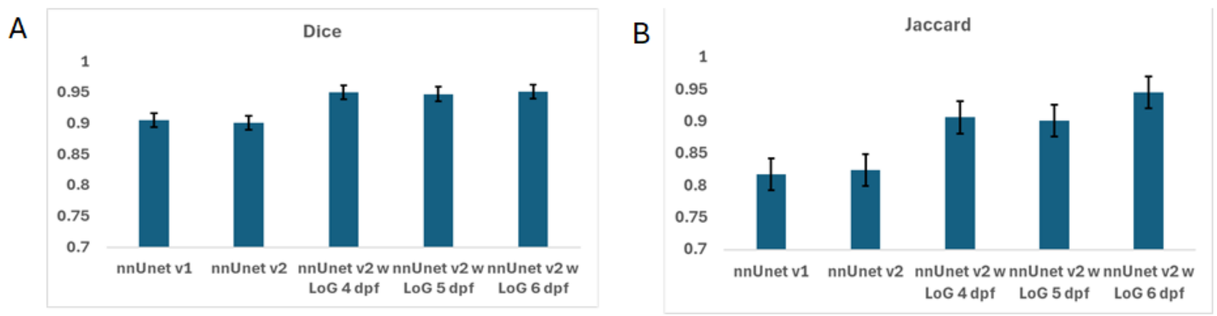


**Supplementary Figure 3. Validation for nnUnet** (A)Dice coefficient scores for varying nnUnet configurations for image preprocessing (B) Jaccard scores for varying nnUnet configurations for image preprocessing
